# The Role of Sleep Duration and Age in Neural Reactivation and Episodic Memory Performance Across the Adult Lifespan^*^

**DOI:** 10.1101/2025.11.24.690290

**Authors:** Masoud Seraji, Soroush Mirjalili, Sir-Lord Wiafe, Audrey Duarte, Vince D. Calhoun

## Abstract

Sleep is essential for episodic memory consolidation, involving the reactivation of neural patterns during encoding and retrieval. This study examined how sleep duration influences encoding-retrieval similarity (ERS) and memory performance across the adult lifespan. Fifty-five adults (18–79 years) participated in a seven-day protocol, combining actigraphy-based sleep monitoring with EEG and memory assessments during a paired associate learning task. Significant correlations between sleep duration and ERS were observed, with clusters showing positive and negative associations across brain regions. In match context trials, ERS in the left frontal region correlated with sleep duration, and age moderated these effects. In mismatch context trials, positive correlations were identified in the left posterior, left frontal, and right posterior regions. These results highlight the role of sleep in modulating neural reactivation patterns during memory retrieval and emphasize the influence of age. The study advances our understanding of how sleep variability impacts memory consolidation and provides a basis for exploring targeted sleep interventions.

**Clinical Relevance:** This study highlights the impact of habitual sleep variability on neural reactivation patterns and memory retrieval across the adult lifespan. The findings emphasize the moderating role of age and suggest that improving sleep duration may enhance episodic memory performance. These insights could guide clinicians in developing personalized sleep interventions to support cognitive health and mitigate age-related memory decline.

## I. Introduction

Sleep plays a crucial role in the consolidation of episodic memory, which involves the retention and integration of past experiences [1]. Experimental studies manipulating sleep conditions—such as variations in retrieval timing (morning vs. evening), sleep deprivation, and the presence of intervening naps—have consistently demonstrated that episodic memory is influenced by sleep across different age groups. Both younger and older adults show sleep-dependent memory effects, although age-related differences in sleep architecture may impact the extent of consolidation [2].

Moreover, polysomnography (PSG) research has identified specific EEG sleep patterns, such as slow-wave sleep (SWS) and sleep spindles, that correlate with memory enhancement in both populations [3]. However, most of these findings stem from highly controlled laboratory settings, which may not fully reflect naturalistic sleep patterns observed in home environments. Additionally, many studies are limited to single-night assessments, necessitating further research to explore long-term memory effects under real-life sleep conditions.

In addition, Poor sleep may contribute to declines in memory function in older adults. Compared to younger individuals, older adults tend to perform worse on memory tasks that require recollection-based retrieval, while their familiarity-based recognition remains relatively preserved [4]. For instance, associative memory performance declines more significantly with age than item recognition [5]. Interestingly, these same types of memory tasks are particularly dependent on sleep in both younger and older adults[6]. Age-related differences in sleep quality have been linked to poorer episodic memory performance, as demonstrated through self-report measures, actigraphy, and polysomnography [7], [8], [9]. Collectively, these studies highlight a consistent relationship between sleep quality and episodic memory performance across different age groups, reinforcing the critical role of sleep in memory consolidation throughout the lifespan.

Actigraphy is a commonly used method for assessing habitual sleep, measuring key parameters such as total sleep time, sleep efficiency, and wake after sleep onset. Neurobiological models of memory propose that successful episodic memory retrieval depends on the reactivation of neural activity patterns from the encoding phase [10]. This concept is supported by encoding-retrieval similarity (ERS) analyses, which show that reactivation of encoding-related neural patterns enhances memory accuracy. Additionally, neuroimaging evidence suggests that event-specific neural reactivation is crucial for successful recollection. While ERS has been linked to episodic memory performance in young adults, the influence of habitual sleep quality on this relationship remains underexplored. This study addresses this gap by collecting one week of habitual sleep data using wrist-worn accelerometers, capturing both average sleep quality and night-to-night variability, to examine their impact on episodic memory performance [11].

In this study, we utilized representational similarity analysis (RSA) to examine time-frequency EEG data recorded during a paired associate learning task. Specifically, we assessed event-specific oscillatory similarities across different frequency bands between encoding and retrieval phases across the adult lifespan. Given the well-established relationship between sleep quality and memory performance in both young and older adults [12], we hypothesized that individual differences in memory performance and ERS would be linked to variations in sleep quality. Additionally, we investigated the effects of age on these associations, aiming to understand how lifespan-related changes influence the relationship between sleep, memory, and neural reactivation patterns.

## II. Methods

### A. Data Collection

The study included 55 right-handed adults (28 females, 25 males, 1 non-binary, and 1 transgender woman), aged 18 to 79 years. Participants were categorized into two groups: young adults (18-36 years) and older adults (56-79 years). All individuals self-identified as native English speakers were right-handed and had normal or corrected-to-normal vision. None reported any uncontrolled psychiatric, neurological, or sleep disorders, nor any history of vascular disease. All participants signed consent forms, and the entire study was approved by the University of Texas at Austin Institutional Review Board.

### B. Experimental Design

The study followed a seven-day experimental protocol, incorporating both laboratory visits and at-home data collection. On day 0, participants attended an initial lab session where they completed a neuropsychological assessment and were fitted with Actiwatch 2 accelerometers (Philips) to monitor habitual sleep patterns. Days 1 and 2 involved at-home sleep tracking, with nightly sleep data recorded remotely. On day 3, participants returned to the lab to complete a memory encoding session followed by an immediate retrieval task. Days 4 to 6 continued with at-home sleep monitoring, culminating in a final lab visit on day 7 for a delayed memory retrieval assessment (Fig. 1). The memory task was designed to assess episodic memory performance. During encoding, participants were shown images of objects placed within scenes, each displayed for four seconds, followed by a 350-750 ms fixation cross. Their task was to evaluate whether the object logically fit within the scene, selecting from three response options: “Yes” (1), “No” (2), or “Somewhat” (3). During retrieval, participants viewed a series of images and categorized them into one of three conditions: “Same Old” (object and background remained unchanged), “Different Old” (the object was the same, but the background had changed), or “New” (entirely novel object) (Fig. 1).

**Figure 1.**
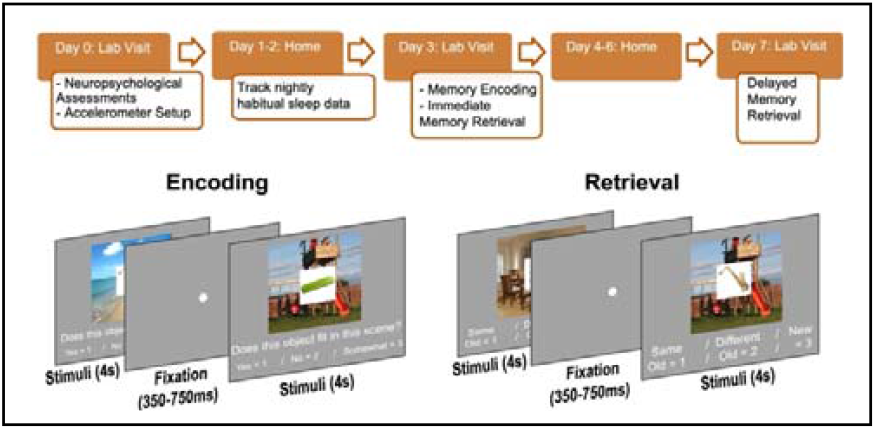
The experimental design involved a combination of laboratory sessions and at-home sleep monitoring using accelerometers. Participants engaged in a memory encoding task where they determined whether an object logically fit within a scene. During the retrieval phase, they categorized the objects as “Same Old” (both the object and background were unchanged), “Different Old” (the object was the same, but the background had changed), or “New” (a novel object). The study spanned seven days, with habitual sleep data continuously collected between memory tasks to assess the relationship between sleep patterns and memory performance.

This structured design allowed for an integration of sleep monitoring and memory performance assessments, facilitating insights into the relationship between habitual sleep patterns and episodic memory consolidation across multiple time points.

### C. EEG Recording and Preprocessing

EEG data were continuously recorded from 31 scalp electrodes using the Brain Vision ActiCAP system, following the extended 10–20 electrode placement system. Electrodes were positioned at Fp1, Fz, F3, F7, FT9, FC5, FC1, C3, T7, TP9, CP5, CP1, Pz, P3, P7, O1, Oz, O2, P4, P8, TP10, CP6, CP2, C4, T8, FT10, FC6, FC2, F4, F8, and Fp2. Additionally, two electrodes at the lateral canthi of the eyes recorded horizontal electrooculogram (HEOG), while two electrodes above and below the right eye captured vertical electrooculogram (VEOG). EEG signals were sampled at 500 Hz with no initial filtering applied.

Offline processing [13], [14], [15] [16], [17] was conducted using MATLAB, incorporating the EEGLAB [18] toolbox. The data underwent baseline correction within the time window of 400 to 200 ms prior to stimulus onset, and the sampling rate was down-sampled from 500 Hz to 250 Hz. Re-referencing was performed using the average of the left and right mastoid electrodes, followed by bandpass filtering between 0.05 and 80 Hz. To eliminate electrical interference, 60 Hz line noise was removed. Independent component analysis (ICA) was then applied to isolate and remove ocular artifacts such as blinks and eye movements. Trials with voltage fluctuations exceeding 150 microvolts were automatically rejected, with an additional manual review conducted to ensure data integrity.

Time-frequency analysis was performed using Morlet wavelet transformations, with 38 frequencies evenly spaced between 3 and 40 Hz. The data were further down-sampled from 250 Hz to 50 Hz, and each trial was segmented to capture the relevant time window from 0 to 2,400 ms post-stimulus onset. This transformation produced a data matrix organized as trials × electrodes × frequency bands. Power estimates were calculated across 120 time intervals, each spanning 20 milliseconds, aligned with the adjusted 50 Hz sampling rate (See [12]). Electrodes were grouped into four non-overlapping regions, and signals from electrodes within each region were averaged. The wavelet-transformed data were then divided into 22 overlapping time windows of 300 ms, with each window advancing by 100 ms [12].

### D. Data Analysis

In this study, RSA was applied to investigate the relationship between EEG activity and memory performance during a visual memory encoding and retrieval task. In the encoding phase, participants were presented with visual scenes and asked to determine whether a specific object logically fit within the scene. During the retrieval phase, they were shown a combination of previously encountered and novel scenes and were required to classify the objects as matching or mismatching those seen during encoding. EEG oscillatory power was recorded across a frequency range of 3 to 40 Hz during both encoding and retrieval. Power values were averaged within 300 ms time windows for each electrode, log-transformed, and subsequently averaged across predefined brain regions (e.g., left frontal cortex). The analysis focused on differentiating between four key memory response types: hits, where participants correctly recognized a previously seen scene as a match; misses, where they failed to recognize a previously seen scene and mistakenly identified it; correct rejections, where they accurately identified an altered scene as mismatched; and false alarms, where they incorrectly classified a mismatched scene as a previously seen match.

To examine neural similarity between encoding and retrieval, Pearson correlations were computed between EEG power vectors across 300 ms time windows. Within-event similarity, comparing retrieval events with their corresponding encoding events, was contrasted with between-event similarity, which compared retrieval events with all encoding events from the same category. These analyses were conducted for hits, misses, correct rejections, and false alarms to assess how neural patterns distinguish successful from unsuccessful memory retrieval.

Next, between-event similarity matrices were subtracted from within-event similarity matrices for each trial (“(1)” and “(2)”), and the resulting values were averaged across trials of the same type for each participant. This produced a time–time similarity matrix for each electrode region and trial condition. To isolate memory performance effects, the average similarity of missed trials was subtracted from that of hit trials, quantifying differences in neural activation patterns associated with successful and unsuccessful memory retrieval.

Specifically, we calculated the difference between within-event and between-event similarities for hits and misses:

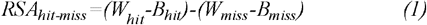

Also, the difference between within-event and between-event similarities for correct rejection (CR) and false alarm (FA):

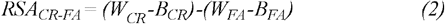

In both formula W is within event similarity, and B is between-event similarity.

### D. Actigraphy

We extracted key sleep metrics, including measures of sleep duration (total sleep time, wake after sleep onset (WASO), and the number of wake bouts) and sleep efficiency (overall efficiency and sleep onset latency). Sleep duration refers to the total time spent asleep during a sleep period, while WASO quantifies the total time spent awake after initially falling asleep. Wake bouts represent the number of times a participant awakened during sleep. Sleep efficiency is defined as the percentage of time spent asleep relative to total time in bed, and sleep onset latency refers to the time taken to transition from wakefulness to sleep. To analyze the relationship between sleep and memory retention, we calculated the average of five sleep metrics following the encoding phase, which we termed the retention period. We then conducted two separate principal component analyses (PCA) to identify key patterns within the data. For our analysis, we selected sleep duration as the focal metric and proceeded with further investigations based on this measure.

### E. atistical Analysis

To identify significant clusters, we examined the correlations between sleep metrics and event-specific encoding-retrieval similarity across encoding periods, aiming to explore the relationship between sleep variability and neural reactivation patterns. Notably, this study incorporated delayed retrieval, providing deeper insights into how neural activity patterns over time influence long-term memory recall. The statistical significance of these temporally clustered correlations was evaluated through a permutation approach, where correlation values were randomly shuffled 10,000 times to generate a null probability distribution for cluster-based statistical comparisons. Morever, to investigate the effect of age on clusters that showed significant associations with sleep duration, we applied a linear regression model. The model included ERS as the dependent variable and age, sleep duration, and their interaction (age * sleep duration) as predictors, expressed as: ERS ∼ Age + Sleep Duration + Age * Sleep Duration.

## III. Results

As outlined in the methods section, we computed correlations between sleep duration and event-specific encoding-retrieval similarity for both match attended context trials (hits and misses) and correct mismatch attended context trials (correct rejections and false alarms).

### A. Match Attended Context Tirals

The clusters where sleep duration demonstrated significant correlations are presented in Fig. 2. For example, in the left frontal region, we identified two distinct clusters with opposing relationships to sleep duration. The first cluster exhibited a significant negative correlation during the encoding window of 300–1200 ms and the retrieval window of 0–800 ms. Conversely, the second cluster displayed a significant positive correlation, spanning the encoding window of 700–1300 ms and the retrieval window of 1100– 1400 ms. In this second cluster, the interaction between age and sleep duration also showed statistical significance, indicating that the effect of sleep duration on ERS is modulated by age.

**Figure 2.**
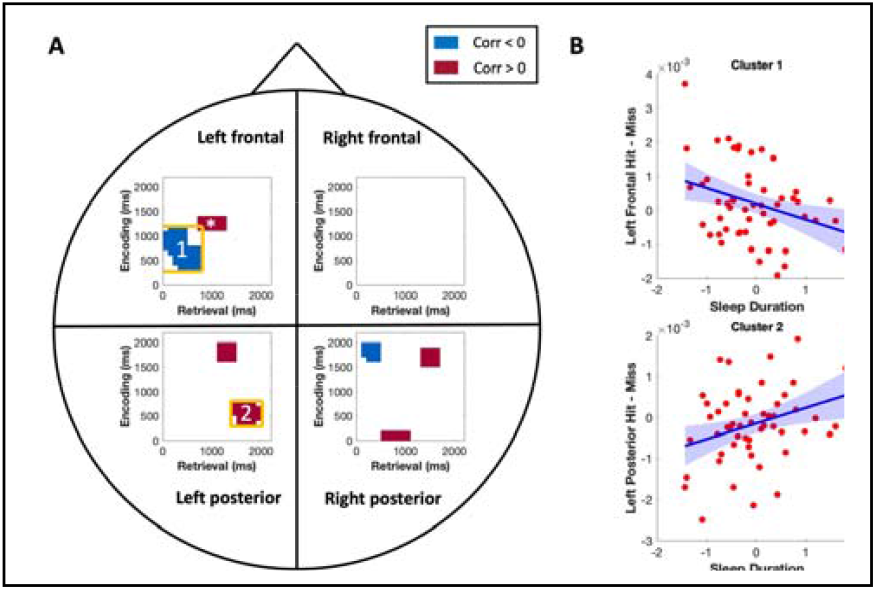
Encoding-retrieval similarity (ERS) time-time clusters for object-scene pairs showing correlations with sleep. (A) Significant time intervals where mean sleep restlessness correlates with event-specific ERS are highlighted. The y-axis represents encoding intervals, and the x-axis represents retrieval intervals (in milliseconds). Clusters with positive correlations are shown in red, while negative correlations are in blue. Statistical significance was determined using a cluster-based permutation method. Clusters marked with a star indicate additional significant effects of age and the interaction between age and sleep. (B) The relationship between event-specific ERS and mean sleep restlessness is illustrated with 95% confidence intervals for two example clusters from (A), one showing a positive correlation and the other a negative correlation, with similar patterns observed in other clusters.

### B. Mismatch Attended Context Tirals

The clusters where sleep duration significantly correlates with event-specific ERS in mismatched context trials are shown in Fig. 3. For example, in the left posterior region, a single cluster demonstrated a significant positive correlation with sleep duration. This cluster spanned an encoding window of 400–1200 ms and a retrieval window of 500–1000 ms. Beyond the left posterior region, additional significant clusters were observed in the left frontal and right posterior regions, all showing positive correlations with sleep duration (Fig. 3). Notably, one cluster in the left frontal region (marked with a star in Fig. 3) indicates that the relationship between sleep duration and ERS is influenced by age, highlighting the role of age as a moderating factor in these neural patterns.

**Figure 3.**
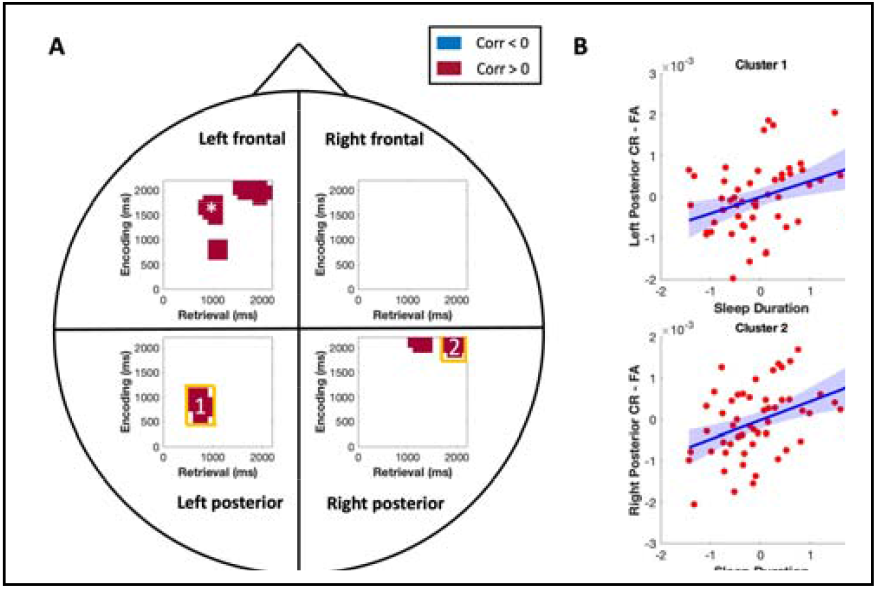
Encoding-retrieval similarity (ERS) clusters showing correlations with sleep duration for correct rejection (CR) and false alarm (FA) trials. (A) Time-time clusters are displayed for four brain regions (left frontal, right frontal, left posterior, and right posterior). The y-axis represents encoding time (ms), and the x-axis represents retrieval time (ms). Clusters with significant positive correlations are shown in red, while blue indicates negative correlations. Clusters marked with a star (*) indicate additional significance for interactions between sleep duration and age. (B) Scatterplots for two example clusters (labeled 1 and 2 in A) illustrate the relationship between sleep duration and ERS values, with blue lines representing regression fits and shaded areas showing 95% confidence intervals.

## IV. Discussion

In this study, we used EEG to examine episodic neural reinstatement effects, focusing on image-pair-specific activity patterns. We observed that neural activity during encoding and retrieval was correlated within similar time intervals, indicating a strong temporal alignment. Interestingly, we also identified significant asymmetries in ERS effects, where encoding activity correlated with retrieval activity occurring either earlier or later in time. These symmetrical and asymmetrical reinstatement effects were influenced by variations in sleep duration, suggesting that sleep may play a critical role in modulating the efficiency and timing of these neural reactivation processes. This modulation potentially impacts how effectively memory traces are accessed and reinstated during retrieval [19]. Our findings extend previous research by showing that sleep not only affects overall memory performance but also influences the temporal dynamics of neural reinstatement critical for successful episodic retrieval.

Clusters in match event trials revealed both positive and negative correlations between ERS and sleep duration. These findings are consistent with earlier research, which similarly identified both positive and negative relationships between sleep metrics and ERS [12], [20]. Furthermore, this study extends our previous work, where we demonstrated both positive and negative clusters between ERS and sleep restlessness [12]. Here, we replicate these effects using sleep duration as a key variable, confirming its influence on neural reinstatement patterns. In contrast, clusters in mismatch event trials showed only positive correlations between ERS and sleep duration. This suggests that longer sleep duration directly enhances the ability to reject false memories in both younger and older adults, highlighting the role of sleep in supporting accurate memory discrimination across age groups [21].

Our findings revealed that the relationship between sleep duration and left frontal ERS in both matched and mismatched trials was moderated by chronological age. This aligns with prior research demonstrating event-specific neural reinstatement over frontal regions in young adults [22]. Furthermore, the late time course of the observed frontal ERS effects is consistent with ERP studies, which have shown sustained frontal ERPs distinguishing recollected events based on study task history or associated perceptual contexts, such as face versus scene contexts [23]. A plausible interpretation of these frontal ERS effects is that they reflect post-retrieval monitoring processes mediated by the prefrontal cortex (PFC). These processes likely include the maintenance, manipulation, and evaluation of retrieved memory representations to support accurate decision-making [24]. These findings emphasize the critical role of frontal neural activity in regulating memory retrieval processes and underscore the influence of sleep duration on these mechanisms, particularly in younger individuals.

In summary, the findings reinforce the link between sleep quality and episodic memory retrieval, highlighting how sleep affects both the occurrence and timing of neural reactivation. These insights could inform interventions to enhance memory, particularly for individuals with sleep disturbances. Future research should explore how specific sleep stages influence ERS and assess sleep-based interventions to improve memory performance, especially in older adults.

## V. Conclusion

This study highlights the significant role of sleep duration in modulating neural mechanisms underlying episodic memory retrieval across the adult lifespan. Using EEG and RSA, we found that sleep duration influences encoding-retrieval similarity patterns, with match trials showing both positive and negative correlations and mismatch trials displaying exclusively positive correlations. These findings suggest that sleep duration enhances the ability to reject false memories and supports effective memory retrieval. The relationship between sleep duration and left frontal ERS was moderated by age, emphasizing lifespan-related differences in memory processes. These results extend previous research by showing that sleep not only affects overall memory performance but also the temporal dynamics of neural reinstatement. The findings underscore the role of the prefrontal cortex in post-retrieval processes such as maintaining and evaluating memory representations. This study provides a foundation for future research to explore the effects of specific sleep stages on memory and to develop sleep-based interventions to improve cognitive function, particularly in older adults or individuals with sleep disturbance.

## Acknowledgment

This research was supported by the National Science Foundation (NSF) through grant numbers 2112455, 2316421, and 2152492.

